# Cultivating efficiency: High-throughput growth analysis of anaerobic bacteria in compact microplate readers

**DOI:** 10.1101/2023.10.10.561742

**Authors:** Oona L.O. Snoeyenbos-West, Christina R Guerrero, Makaela Valencia, Paul Carini

## Abstract

Anaerobic microbes play crucial roles in environmental processes, industry, and human health. Traditional methods for monitoring the growth of anaerobes, including plate counts or subsampling broth cultures for optical density measurements, are time and resource intensive. The advent of microplate readers revolutionized bacterial growth studies by enabling high-throughput and real-time monitoring of microbial growth kinetics but their use in anaerobic microbiology has remained limited. Here, we present a workflow for using small-footprint microplate readers and the Growthcurver R package to analyze the kinetic growth metrics of anaerobic bacteria. We benchmarked the small-footprint Cerillo Stratus microplate reader against a BioTek Synergy HTX microplate reader in aerobic conditions using *Escherichia coli* DSM 28618 cultures. The growth rates and carrying capacities obtained from the two readers were statistically indistinguishable. However, the area under the logistic curve was significantly higher in cultures monitored by the Stratus reader. We used the Stratus to quantify the growth responses of anaerobically grown *E. coli* and *Clostridium bolteae* DSM 29485 to different doses of the toxin sodium arsenite. The growth of *E. coli* and *C. bolteae* was sensitive to arsenite doses of 1.3 μM and 0.4 μM, respectively. Complete inhibition of growth was achieved at 38 μM arsenite for *C. bolteae*, and 338 μM in *E. coli*. These results show that the Stratus performs similarly to a leading brand of microplate reader and can be reliably used in anaerobic conditions. We discuss the advantages of the small format microplate readers and our experiences with the Stratus.

**Importance statement:** We present a workflow that facilitates the production and analysis of growth curves for anaerobic microbes using small-footprint microplate readers and an R script. This workflow is a cost and space-effective solution to most high-throughput solutions for collecting growth data from anaerobic microbes. This technology can be used for applications in which high-throughput would advance discovery, including microbial isolation, bioprospecting, co-culturing, host-microbe interactions, and drug/toxin-microbial interactions.

## INTRODUCTION

Measuring microbial growth is crucial for understanding the behavior and physiology of single-celled organisms (Hall et al. 2014; Monod 1949; Koch 1961). Optical density (OD) measurements are commonly used to monitor microbial growth, and with the advent of automated microplate readers and software packages, the collection of OD data has become more efficient (Kurokawa and Ying 2017; Casagrande Pierantoni et al. 2019; Atolia et al. 2020).

Microplate readers allow many samples to be analyzed simultaneously, providing reliable and efficient measurements of growth rates and other kinetic data. This has significantly improved the accuracy and speed of high-throughput data collection and analysis in microbiology research. For example, microplate readers have been used to study the growth responses to antibiotics or environmental toxins (Krishnamurthi et al. 2021; Bollenbach et al. 2009; Li et al. 2016; Begot et al. 1996), the chronological life span in yeast (Jung et al. 2015), metabolic footprinting in *Pseudomonas* (Pedersen et al. 2021), and fitness effects in long-term evolution experiments (Atolia et al. 2020), among other applications.

Relative to aerobic microbes, tracking the growth dynamics of anaerobic microbes adds a layer of complexity due to the need for culture manipulations and incubations to be conducted without exposure to oxygen. Researchers typically culture anaerobic microbes in Hungate or Balch-type culture tubes with thick rubber septa (Börner 2016). Unfortunately, there are no automated tools currently available to measure optical density from cultures incubated in these types of tubes (Vuono et al. 2019), meaning researchers must manually pierce the rubber septa with a needle to extract a small volume of culture to be read by a spectrophotometer. This process is labor-intensive, low throughput, and relies heavily on the efficiency of the researcher conducting the experiments.

Several approaches have been used to circumvent the low-throughput nature of anaerobe cultivation. One approach excluded oxygen by sealing lids onto microtiter plates and flushing the headspace with N_2_ to form anaerobic conditions (Koutny and Zaoralkova 2005). A second approach enzymatically removed oxygen from the culture medium ahead of growth analysis (Lam et al. 2018). These two approaches attempt to limit oxygen exposure while the plate is incubated outside of an anaerobic chamber in oxygenated conditions—an approach that may not work well for all microbes. In contrast, the environment of the microplate reader itself can be modified by relocating it into an anaerobic chamber (Luna et al. 2022; Candry et al. 2018; Aranda-Díaz et al. 2022; Zünd et al. 2023). However, many microplate readers are too large to fit through anaerobic chamber airlocks and require connections to an external computer for data collection. Anaerobic chambers are also typically restrictive, providing just enough room for researchers to carry out necessary manipulations and incubations. Additionally, microplate readers are expensive instruments that are financially inaccessible to many laboratories. Because of the high cost, they are often purchased as shared laboratory equipment or are placed in core facilities, where they need to be accessible for a wide variety of applications. Thus, the substantial physical size, the common need for connection to a computer, and the cost of many microplate readers have been obstacles to their widespread adoption in anaerobic chambers (Jensen et al. 2015)

Here, we present a high throughput cultivation and growth analysis pipeline based on inexpensive small-footprint microplate readers to overcome some of the challenges associated with microbial growth analysis in anaerobic conditions (Fig. 1). These small-format microplate readers—together with a small-footprint shaker—facilitate the collection of OD_600_ measurements from cultures growing in an anaerobic chamber. We show that the mean growth rates and carrying capacities derived from modeling OD_600_ measurements of aerobic bacteria collected with the small-footprint reader did not vary significantly from those rates collected and analyzed with a state-of-the-art popular brand of microplate reader. We used the small footprint microplate reader to quantify the effect of arsenite exposure on the anaerobic growth of two bacterial strains belonging to the mouse intestinal bacterial collection (miBC; Lagkouvardos et al. 2016). Our approach is translatable to high throughput growth studies on anaerobic microbes from any environment.

**Figure 1.**
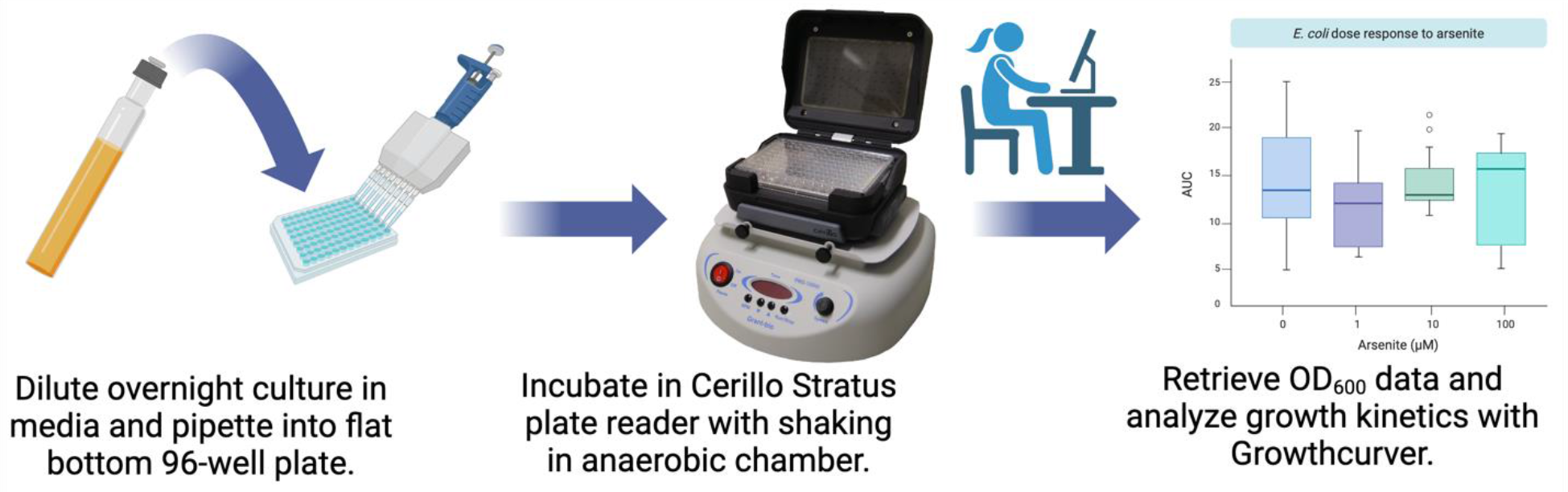
Conceptual diagram of small-footprint plate reader workflow. Figure constructed with Biorender.com.

## RESULTS & DISCUSSION

We sought to develop a cost-effective high-throughput strategy to measure and model the growth of anaerobic bacteria (Fig. 1). This workflow is based on the Cerillo Stratus microplate reader, which captures OD_600_ and temperature measurements at user-defined time points and stores the data in comma separated value (.csv) or text (.txt) files on a MicroSD card. The output data contains measurement timestamps, timepoint temperature, and OD_600_ readings for each well of a 96-well microplate. We paired the Stratus microplate reader with a Grant Instruments Microplate Shaker PMS-1000i small-footprint orbital shaker. This shaker was chosen because it fits inside of COY Model 2000, Forced Air Incubator for Vinyl Chambers that are commonly sold with COY anaerobic chambers. We modeled microbial growth using custom scripts and Growthcurver in R (Sprouffske and Wagner 2016).

We benchmarked the efficacy of the Stratus to obtain accurate growth curve data by comparing the growth rates obtained from the Stratus microplate reader to those obtained from a BioTek Synergy HTX that is commonly used to monitor microbial growth. To do this, we grew aerobic overnight cultures of *Escherichia coli* DSM 28618 and used them to inoculate two 96-well microtiter plates containing Lysogeny Broth (LB) media. We inoculated 64 wells and left 32 wells uninoculated. The plates were covered with Breathe-Easy® Sealing Membranes to facilitate gas exchange and incubated in either the Synergy HTX or the Stratus reader. The plate in the Synergy HTX microplate reader was incubated at 37ºC with continuous shaking at 180 RPM using Synergy HTX’s internal temperature control and shaking functionality. The microplate incubated in the Stratus was affixed to a PMS-1000i microplate shaker, shaking at 180 RPM inside of a 37ºC incubator. Both microplate readers were operating outside of an anaerobic chamber in aerobic conditions. The OD_600_ was measured every 3 minutes overnight (∼18-20 h).

We downloaded the OD_600_ data from both microplate readers and modeled the resulting kinetic parameters with the Growthcurver R package (Sprouffske and Wagner 2016).

Growth rates and carrying capacities from the Stratus and Synergy HTX microplate readers did not differ significantly, but the area under the curve displayed significant differences across the two readers (Fig. 2). The mean carrying capacities (*k* in Growthcurver) obtained from the two readers were statistically indistinguishable (Fig. 2a; Mann-Whitney U *p*=0.088). The mean carrying capacity obtained from the Stratus was 0.51 ± 0.02 (mean ± SEM; n=64) compared to 0.47 ± 0.01 (mean ± SEM; n=64) for the Synergy HTX (Fig. 2a). Similarly, the mean growth rates (*r* in Growthcurver) across the two devices did not differ significantly (Fig. 2b; Mann-Whitney U *p*=0.094). The average growth rate of *E. coli* cultures incubated in the Stratus microplate reader was 1.24 ± 0.06 h^-1^ (mean ± SEM; n=64) compared to 1.28 ± 0.02 h^-1^ (mean ± SEM; n=64) for cultures grown in the Synergy HTX. Growthcurver calculates a metric that integrates growth information from the logistic parameters derived from *r* and *k* called *area under the curve* (AUC; *auc_l* in Growthcurver). The AUC values derived from the Stratus (10.2 ± 0.36 mean ± SEM; n=64) were significantly higher than those obtained from the Synergy HTX (7.65 ± 0.21 mean ± SEM; n=64) (Fig. 2c; Mann-Whitney U *p*=1.14e-14).

**Figure 2:**
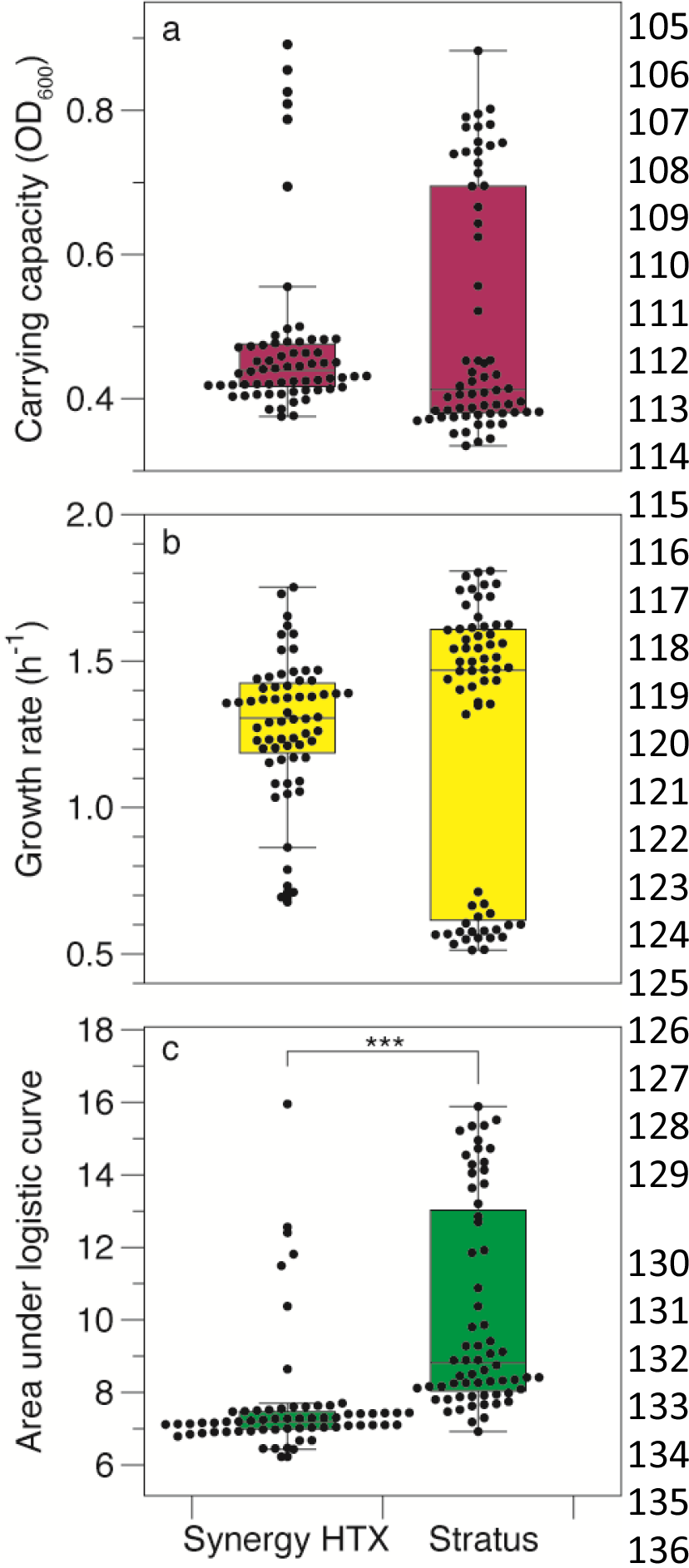
Stratus plate readers generated mean growth rates and maximum carrying capacities indistinguishable from the Synergy HTX. Points are the kinetic parameters calculated from microbial growth in each inoculated well, including (a) carrying capacity; (b) growth rate; and (c) area under the logistic curve. All values were calculated from OD_600_ measurements by the respective microplate reader using Growthcurver. Each point represents a measurement from a single replicate well. Box plots illustrate interquartile range ± 1.5 × interquartile range. The horizontal line in each box plot is the median. Outliers (>1.5 × interquartile range) are shown as points. The bracket with asterisks in (c) denotes a significant difference (Mann-Whitney U p-value ≤0.001)

The growth data collected from the Stratus plate was bimodally distributed for all three growth parameters (Fig 2). Inspection of the growth curves showed a growth curve variant that was characterized by a higher carrying capacity and a slower growth rate resulting in a higher AUC (Supplementary Figures 1 & 2). This growth variant was identified in both the Synergy HTX and the Stratus-incubated microtiter plates (Supplementary Figures 1 & 2, respectively). However, the proportion of wells displaying this variant was higher in the Stratus (33% of wells) than in the Synergy HTX (8% of wells). Previous work in *E. coli* showed growth rate bimodality was dependent on the initial density of cells in each well (Irwin et al. 2010). Yet, in our experiments, both plates were inoculated from the same culture suspension, making significant differences in initial cell densities an unlikely explanation. We speculate that mixing differences between the two shakers—either in actual shaking rate or the orbit diameter—may explain the observed differences. Subtle differences in shaking may have affected the oxygenation of the cultures, leading to the observed growth differences.

After confirming the efficacy of the Stratus microplate reader in aerobic conditions, we used it in anaerobic conditions. We set up an experiment to quantify how the AUC of *E. coli* DSM 28618 and *Clostridium bolteae* DSM 29485 responded to different doses of arsenite when growing anaerobically. AUC has been used previously to quantify the degree of inhibition in microbial cultures (Tiina and Sandholm 1989; Lambert and Pearson 2000). We initiated overnight cultures of *E. coli* or *C. bolteae* from freezer stocks and used the culture to anaerobically inoculate 200 μl cultures in polystyrene microwell plates containing the appropriate growth medium. We added arsenite at eight doses: 0, 0.4, 1.3, 3.8, 38.5, 116, 335, or 1,039 μM with 8 replicates per dose. The plates were covered with Breathe-Easy® Sealing Membranes to allow gas exchange and incubated with shaking at 180 RPM at 37ºC overnight in Stratus microplate readers inside a COY anaerobic chamber with an 95%:5% N_2:_H_2_ atmosphere.

The mean AUC for anaerobically grown *C. bolteae* and *E. coli* was significantly variable across the range of concentrations applied (Fig. 3, Kruskal-Wallace p-value 1.43×10^-9^ and 3.48×10^-8^, respectively). *C. bolteae* grown in the absence of arsenite had a mean AUC of 14.9 ± 3.04 (mean ± SD). AUC was reduced at arsenite doses of 0.4-3.8 μM, relative to no arsenite controls, but not significantly (Fig. 3a, Dunn’s Bonferroni-corrected *p*-value ≥0.05). We defined complete growth inhibition as treatments with modeled AUC that were statistically indistinguishable from AUC calculated from the uninoculated wells. Complete inhibition of *C. bolteae* was observed at arsenite concentrations greater than 38 μM (Fig. 3a; Dunn’s Bonferroni-corrected *p*-value ≥0.05 relative to uninoculated blanks). In contrast, *E. coli* displayed a mean AUC of 3.33 ± 0.25 and 0.4 μM arsenite had no significant effect on the AUC (Fig. 3b; Dunn’s Bonferroni-corrected *p*-value >0.05). Arsenite doses of 1.3-116 μM reduced growth, albeit not significantly compared to no arsenite treatments (Fig. 3b, Dunn’s Bonferroni-corrected *p*-value ≥0.05). Arsenite doses of 335 and 1039 μM significantly inhibited growth relative to unamended cultures but were significantly higher than rates modeled from the uninoculated wells (Fig. 3b; Dunn’s Bonferroni-corrected *p*-value ≤0.05). We did not observe bimodality in the AUC under anaerobic conditions. These results are consistent with previous studies on the inhibitory effect of arsenic on bacterial growth. For example, the growth of bacterial isolates from soil and water from the Hazaribagh tannery industrial area in Bangladesh decreased with increasing concentrations of As (Prosun et al. 2020). Goswami et al. (2014) modeled the effect of As (arsenic trioxide, As_2_O_3_) on the growth of *Aeromonas hydrophila* and observed no significant change in bacterial growth within the range of 0-30.32 μM As.

**Figure 3:**
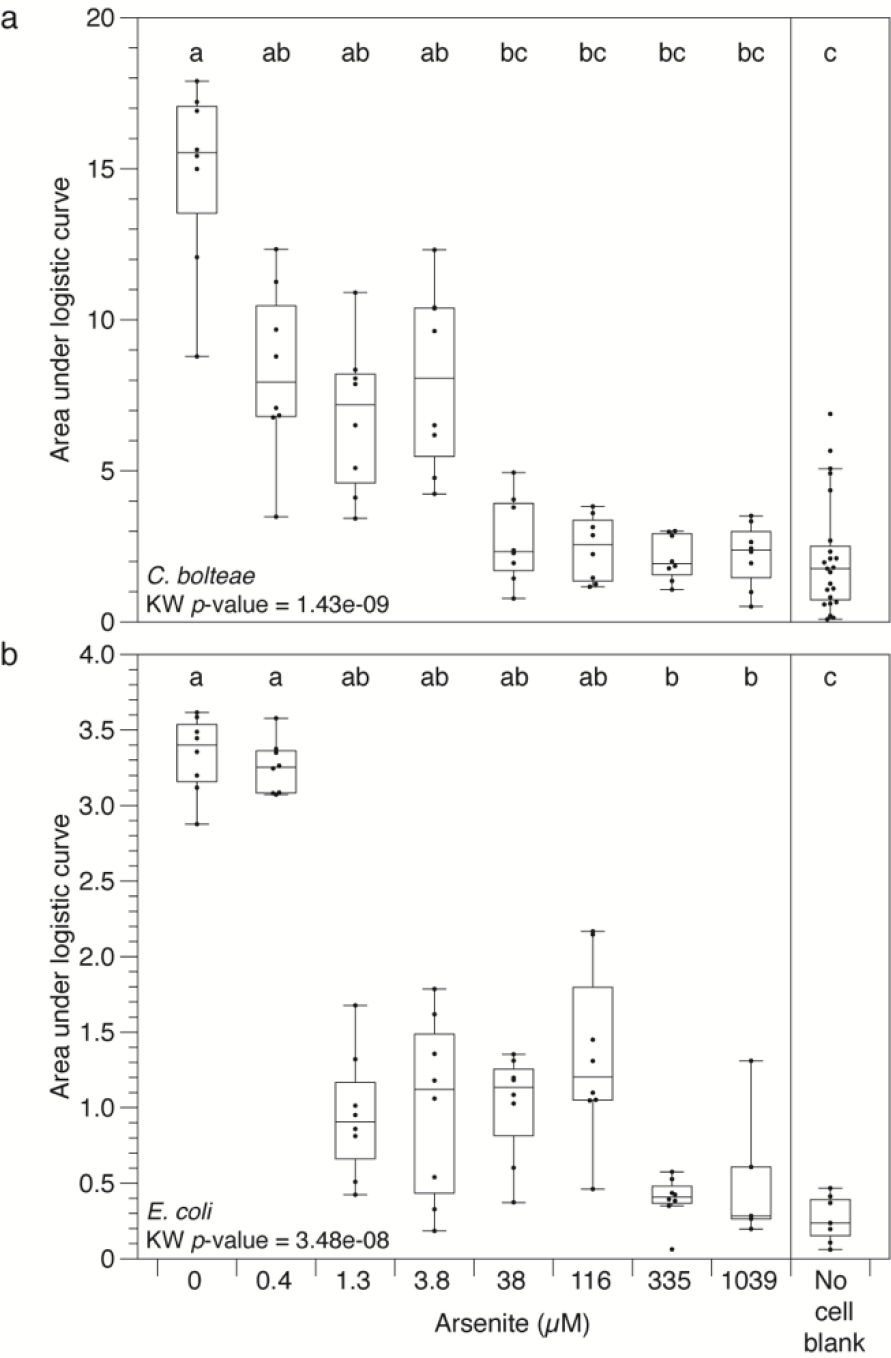
Kinetic responses of (a) C. bolteae and (b) E. coli to arsenite while growing anaerobically. Points are the logistic area under the curve (AUC) derived from growth curves obtained in microtiter plates incubated in the in the Stratus microplate reader. Box plots illustrate interquartile range ± 1.5 × interquartile range. The horizontal line in each box plot is the median. Outliers (>1.5 × interquartile range) are shown as points. Box plots sharing letters are statistically indistinguishable by Dunn’s test as defined by having a Bonferroni-corrected p-value of ≥0.05. KW=Kruskal Wallis.

### Considerations and Difficulties

The Stratus microplate readers generally performed as expected in the anaerobic chamber. Yet, there were areas that we felt the Cerillo under-performed, potentially limiting their utility in certain applications. First, the cultures were continuously shaken in anaerobic conditions during the duration of data collection. However, some anaerobes may be inhibited by agitation (Mustafa, Chua, and El-Enshasy 2019; Dannenberg, Wudler, and Conrad 1997). We did not test the units in the absence of shaking, or with periodic shaking. Importantly, the plate readers and shaking platforms are separate units and there is no integrated method to systematically agitate the plate contents before obtaining time points. Second, while conducting these experiments, we found the Stratus software and/or hardware to be inconsistent. We experienced connection issues and periodic changes or bugs in the format of the data output.

Some of these problems have been addressed with software updates from Cerillo. Occasionally the format of the data collected and stored on the .csv would change (usually coincident with a software update) without advance notification, requiring us to recognize anomalies in our data, track down the change, and adjust our code accordingly. Third, handling the MicroSD card in the anaerobic chamber was not possible because of the small size of the MicroSD card and the reduced dexterity when using anaerobic chamber gloves. Thus, to retrieve the MicroSD card the Stratus unit needed to be removed from the chamber, as there is no built-in wireless data transfer capability. Cerillo has since introduced a wireless accessory package allowing wireless access.

However, this utility is only available with the purchase of their Software as a Service premium software. Finally, several physical constraints of the units as operated in anaerobic chambers should be considered. The Stratus is powered by a USB-C power adapter. The PMS-1000i microplate shaker requires its own power cord. Thus, if the plate is shaken, two outlets are required per plate and reader combination. However, a solitary USB power hub could substitute for electrical outlets for the Stratus readers. Although the Stratus readers are designed to be stackable, we did not attempt to stack multiple Stratus readers on a single shaker and obtain growth readings, but stacking up to two Stratus readers on a single shaker may be possible.

## Conclusion

We present a workflow that relies on small-footprint microplate readers and an R script to analyze the resulting OD_600_ data collected in anaerobic conditions. This approach provided results consistent with a name-brand microplate reader system. The small footprint saves substantial space in anaerobic chambers. We estimate that two Stratus-shaker systems fit within the same footprint as a single Synergy HTX. However, alternative shaker configurations may allow several Stratus instruments to be run simultaneously in the same footprint as the Synergy HTX. Although the small footprint readers are more affordable than the larger systems, they are also more limited in what they can measure (OD at a single preset wavelength). Our pipeline facilitates the quick analysis of many samples inside an anaerobic chamber, increasing the speed of research and providing insight into anaerobe growth and metabolism. This method may be useful for the ever-expanding field of gut microbiome research where high throughput cultivation will spur the discovery of novel metabolites and probiotics and facilitate dose-response studies of various drugs and toxic compounds on the microbiome.

## METHODS & EXPERIMENTAL DESIGN

### Bacterial cultures and media

Bacterial strains *Escherichia coli* DSM 28618 and *Clostridium bolteae* DSM 29485 were purchased from the German Collection of Microorganisms and Cell Cultures (DSMZ, Germany), and grown in the recommended media and incubation temperatures.

*E. coli* was grown in Lysogeny broth (LB) (per liter, 10 g tryptone, 5 g yeast extract, 10 g sodium chloride and adjusted to pH 7 with NaOH). *C. bolteae* was grown in Wilkins Chalgren Anaerobic Broth (CMO643 Oxoid Ltd). Media for anaerobic strains were prepared by boiling and cooling to room temperature while sparging under a constant 100% N_2_ gas flow. All anaerobic media contained 0.5 ml L^-1^ Na-resazurin solution (0.1% w/v) and L-cysteine-HCl (0.3g L^-1^). After autoclaving, media were equilibrated in an anaerobic chamber overnight. All anaerobic experiments were performed at 37°C inside a COY Model 2000, Forced Air Incubator for Vinyl Chambers, in a vinyl anaerobic chamber (Coy Laboratory Products, Inc., Grass Lake, MI, USA). All microplate-incubated cultures were conducted in 200 μl volumes in flat-bottomed polystyrene 96-well plates sealed with Breath-Easy Sealing Membranes and lids. All growth curves were initiated with a 1:100 dilution of overnight cultures pre-incubated for ∼18-20 h under aerobic or anaerobic conditions depending on the experiment.

### Aerobic and anaerobic OD data collection and experimental design

The plate in the Synergy HTX microplate reader was incubated inside the Synergy HTX at 37ºC with continuous shaking at 180 RPM. The microplate incubated in the Stratus was affixed to a PMS-1000i microplate shaker, shaking at 180 RPM inside of a 37ºC incubator. Both microplate readers were operating outside of an anaerobic chamber in aerobic conditions. The OD_600_ was measured every 3 minutes overnight (∼18-24 h).

For the arsenite dose experiments, individual treatments contained no (0 μM), or incremental micromolar doses of the cellular toxin As, as sodium arsenite (NaAsO_2_): 0.4, 1.3, 3.8, 38.5, 115.5, 346, and 1,039 μM. We inoculated 8 replicates per dose. Growth was monitored with an OD_600_ reading taken every 3 minutes in anaerobic conditions at 37°C in a Stratus microplate reader (Cerillo, Charlottesville, VA, USA) shaking on a Microplate Shaker PMS-1000i (Grant Instruments, Shepreth, Cambridgshire, UK) at 180 rpm.

### Growth curve modeling and kinetic parameter estimation

The OD_600_ data was downloaded from the respective microplate readers and modeled with the Growthcurver R package (Sprouffske and Wagner 2016). The Stratus output data was restructured using a custom R script to facilitate Growthcurver analysis.

## Data Processing and Code Availability

After processing the data with Growthcurver, we removed wells that did not model or for which the model was questionable as explained in Growthcurver literature. We also removed data from uninoculated blank wells that displayed growth due to carryover or contamination. All code used is available at: https://github.com/cbaughan/CerilloWrangling.

## Statistical Analysis

All statistical analyses, including Kruskal-Wallis, Mann-Whitney U, and Dunn’s tests were conducted in R.

## ACKNOWLEDGEMENTS

We thank Deanna Sanchez at the BIO5 Institute for the photograph of the Stratus Plate reader included in Figure 1. We also thank Dr. Pawel Kiala for ordering the DSMZ cultures used in this study. Research reported in this publication was supported by the National Institute of Environmental Health Sciences of the National Institute of Health under Award Number P42 ES004940. The content is solely the responsibility of the authors and does not necessarily represent the official views of the National Institute of Health.

## Notes

### Competing Interest Statement

The authors have declared no competing interest.

