## Supplemental Figures 1 & 2 for "Cultivating efficiency: High-throughput growth analysis of anaerobic bacteria in compact microplate readers"

Supplementary Figure 1: Growth curves of *E. coli* incubated in aerobic conditions in a BioTek Synergy HTX microplate reader. Columns 1-8 were inoculated with cells. Columns 9-12 were uninoculated media blanks. The black lines are plotted from OD600 measurements collected by the microplate reader. The overlaying red lines are best-fit curves modeled from Growthcurver. Curves inside of blue boxes are growth curve variants that had a high carrying capacity and a lower growth rate, leading to a higher AUC, relative to the remaining wells.

Supplementary Figure 2: Growth curves of *E. coli* incubated in aerobic conditions in a Cerillo Stratus microplate reader. Columns 1-8 were inoculated with cells. Columns 9-12 were uninoculated media blanks. The black lines are plotted from OD600 measurements collected by the microplate reader. The overlaying red lines are best-fit curves modeled from Growthcurver. Curves inside of blue boxes are growth curve variants that had a high carrying capacity and a lower growth rate, leading to a higher AUC, relative to the remaining wells.
